# Insights into gene expression changes under conditions that facilitate horizontal gene transfer (mating) of a model Archaeon

**DOI:** 10.1101/2020.07.06.189845

**Authors:** Andrea M. Makkay, Artemis S. Louyakis, Nikhil Ram-Mohan, Uri Gophna, J. Peter Gogarten, R. Thane Papke

## Abstract

Horizontal gene transfer is a means by which bacteria, archaea, and eukaryotes are able to trade DNA within and between species. While there are a variety of mechanisms through which this genetic exchange can take place, one means prevalent in the archaeon *Haloferax volcanii* involves the transient formation of cytoplasmic bridges between cells and is referred to as mating. This process can result in the exchange of very large fragments of DNA between the participating cells. Genes governing the process of mating, including triggers to initiate mating, mechanisms of cell fusion, and DNA exchange, have yet to be characterized. We used a transcriptomic approach to gain a more detailed knowledge of how mating might transpire. By examining the differential expression of genes expressed in cells harvested from mating conditions on a filter over time and comparing them to those expressed in a shaking culture, we were able to identify genes and pathways potentially associated with mating. These analyses provide new insights into both the mechanisms and barriers of mating in *Hfx. volcanii*.

## 1. Introduction

In eukaryotic sexual reproduction, the recombination of DNA is intertwined through the processes of meiosis and syngamy, but in prokaryotes the two processes are separated. Prokaryotes reproduce asexually and recombine fragments of DNA between cells in a donor-recipient fashion in a process called horizontal or lateral gene transfer (HGT or LGT) ^1, 2^. There is a diversity of known mechanisms by which DNA can move between cells. The best-characterized means of HGT include transduction, the movement of DNA between cells via a phage or virus; transformation, the uptake of naked environmental DNA (eDNA); and conjugation, the movement of genetic material, usually plasmids (selfish genetic elements), via a narrow channel that transfers single-stranded DNA ^3^.

Very little knowledge exists regarding HGT mechanisms in archaeal species. Natural transformation has been detected in only four archaeal species ^4-7^. However, the molecular machinery by which DNA is transported across the membrane remains elusive. For instance, the required pore protein gene *comEC* and its homologs have not been identified in Archaea ^8, 9^. A cell-cell contact mechanism, the “crenarchaeal system for exchange of DNA” (Ced) ^10^ found in *Sulfolobus acidocaldarius*, which imports DNA from the donor rather than exports it as in bacterial conjugation, requires protein families that have only been identified in crenarchaeal organisms to date ^10^. The oldest report for HGT in an archaeon is the cell-cell contact dependent mechanism called “mating” found in the hypersaline adapted euryarchaeon *Haloferax volcanii* ^11-13^. The mating mechanism exhibited by *Hfx. volcanii* involves fusion of (at least) two cells that results in transfer of DNA and generates a heteroploid state ^14, 15^.

Analysis of mating during biofilm formation demonstrated approximately the same mating efficiency as seen under filter-mediated laboratory experiments, suggesting that biofilms are a natural condition under which *Hfx. volcanii* undergoes HGT, and that biofilm formation and HGT could be intertwined physiological pathways ^16^.

Successful mating has been observed only within or between *Haloferax* species ^17^, probably due to the presence of a self/non-self recognition system. Laboratory experiments tested this hypothesis by measuring the frequency of fusion events (the first step in mating occurring prior to DNA transfer and recombination) within and between *Haloferax* species. Results showed there was a higher efficiency for fusion events within *Haloferax* species, as opposed to between species, by a factor of about 8 ^15^. This self/other distinction may be the result of differential glycosylation of S-layer proteins, as S-layer glycosylation is required for mating ^18,19^. Glycosylation pathways can be highly diverse within haloarchaeal species. *Hfx. volcanii* DS2 expresses different S-layer glycosylation when required to grow at lower-than-optimal salinity ^20^, and these pathways are constructed uniquely amongst individual *Hfx. volcanii* strains that are identical for their *rpoB* gene ^21^. Such variance of S-layer glycosylation patterns implies an impact on self-recognition even within species.

Similar to observations in bacteria, reduced recombination occurs with increased phylogenetic distance in Haloarchaea ^22^. However, Naor et al ^15^ suggest that halophilic archaea are particularly promiscuous in comparison to their bacterial cousins with respect to transfer within and between closely related species. Therefore, the impact of HGT on Haloarchaea is perhaps greater than for other organisms. Notably, Haloarchaea have acquired several hundreds of genes and gene fragments from Bacteria ^23-25^.

To uncover more details about the HGT mating mechanism in *Hfx. volcanii*, a transcriptome approach was used to determine which genes are differentially expressed when moving a population of cells from a shaking culture environment to being immobilized under filter-mating conditions over a 24-hour period. Here the changes in the expression of some groups of genes are examined hypothesized *a priori* to be relevant for the overall mating success in *Hfx. volcanii*. Genes with an expression pattern corresponding to the expectation for genes involved in mating were identified. While further work is needed to unravel the intricacies of the mating process, these data provide a foundation upon which to direct additional research.

### Methods

*Haloferax volcanii* strains H53 (Δ*pyrE2*, Δ*trpA*) and H98 (Δ*pyrE2*, Δ*hdrB*), which are auxotrophs for uracil/tryptophan and uracil/thymidine respectively, were grown at 42°C with shaking in casamino acids (CA) medium ^26^. CA medium contains per liter: 144 g of NaCl, 21 g of MgSO_4_ · 7H_2_O, 18 g of MgCl_2_ · 6H_2_O, 4.2 g of KCl, 5.0 g casamino acids, and 12 mM Tris HCl (pH 7.5). As needed, the medium was supplemented with 50 µg/mL uracil (ura), 50 µg/mL tryptophan (trp), and/or 40 µg/mL both thymidine and hypoxanthine (thy). H53 cultures were grown in CA with uracil and tryptophan and H98 cells were grown in CA with uracil, thymidine, and hypoxanthine.

Liquid cultures of *Hfx. volcanii* were grown until late exponential phase; the cells were then pelleted and resuspended in higher volumes of fresh CA medium (+ura/trp or +ura/thy) and allowed to grow overnight to ensure cells were in mid-log phase when mating. Cells were diluted in CA medium to OD_600_=0.25 and equal volumes of each cell line were mixed together per mating replicate. In the case of suspended (non-mating) cells, the cell cultures were mixed, pelleted, and resuspended directly in either 1 mL Trizol reagent for RNA extraction or unsupplemented CA medium for plating. In the case of mated replicates, 25 mL cell suspension were pushed with a syringe on to 25 mm diameter, 0.45 µm nitrocellulose filters. The filters were subsequently laid, cells up, on plates of solid CA medium (+ ura, trp, thy) for the duration of mating. For each time point (0, 2, 4, 8, and 24 hours), three filters were prepared. At the time of harvesting, cells from one filter were resuspended in 1 mL unsupplemented CA medium by incubating at 42°C for one hour with agitation for plating and the remaining two filters were resuspended in 1 mL Trizol reagent for RNA extraction. The overall mating was done with two separate isolates on three filters at each time point (Fig. 1A; Supplementary Table S1).

**Figure 1:**
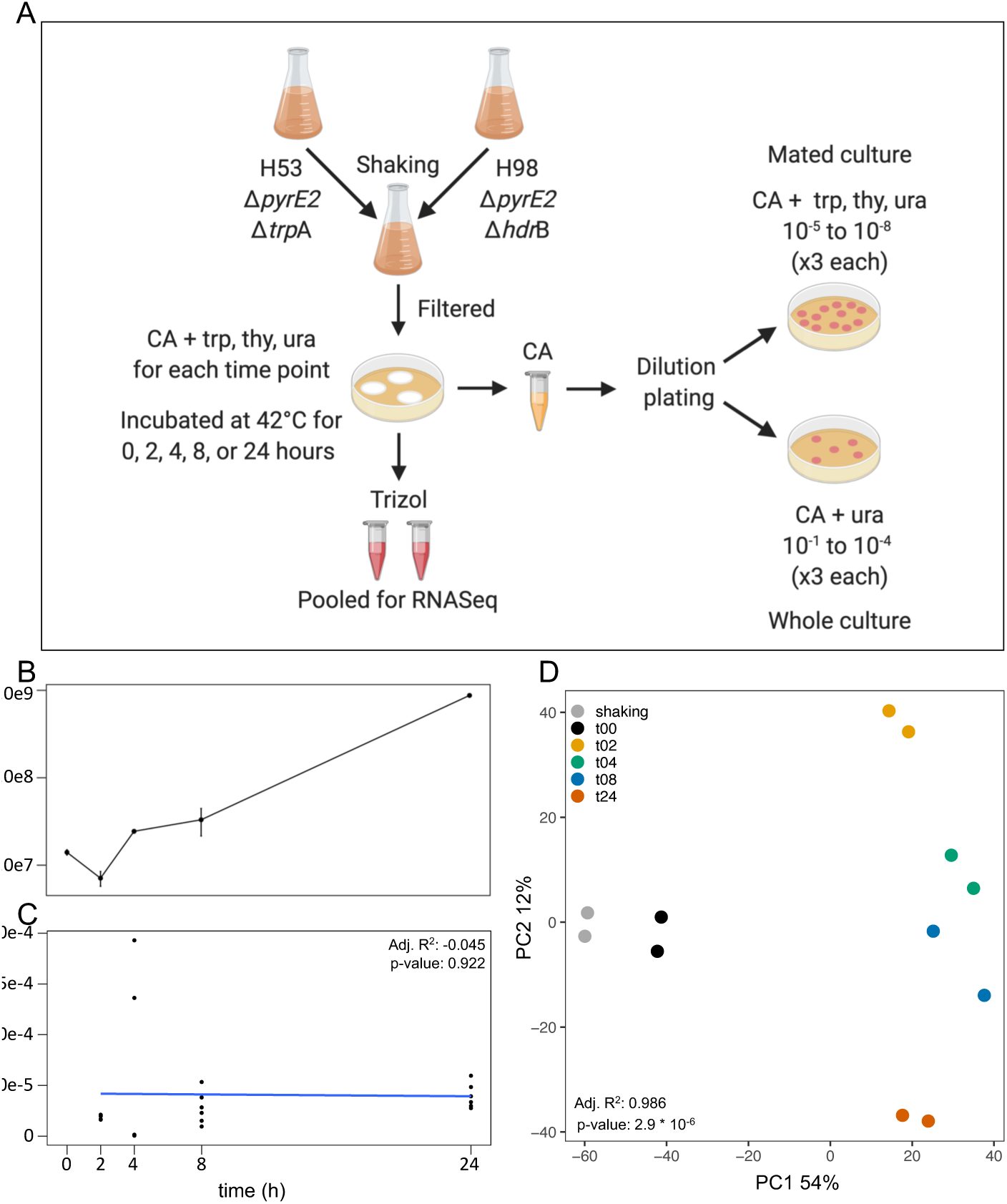
(A) Experimental overview (repeated for two biological replicates): auxotrophic cells were combined and filter plated for mating in casamino acids (CA) medium with uracil (ura), tryptophan (trp), and both thymidine and hypoxanthine (thy); two filters were used for sequencing and one for dilution plating on selective and non-selective media to calculate mean colony forming units (cfu) and mating efficiency; (B) Growth curve of *Haloferax volcanii* plotting mean cfu on log_10_ scale (standard error bars based on triplicate plates for two replicates; raw data are in Supplementary Table S1); (C) mating efficiency based on triplicate plates for two replicates excluding shaking and time 0, linear model regression fit calculated for time 2 to 24 h and comparison tested with ANOVA (n=6 for each time point; raw data are in Supplementary Table S1); and (D) principle component analysis of mating transcriptome time-points identified by replicate pairs and ANOVA tested on PC1 for time.

Mating efficiency was determined by plating cells on selective media. Whole cell counts were determined by plating 10-fold dilutions of cells on solid CA medium supplemented with ura, trp, and thy and mated cell counts were determined by plating dilutions on CA agar supplemented only with ura. Growth on CA + ura does not occur with the auxotrophs H53 or H98, but mating leads to recombination and overcomes the requirement for both trp and thy supplementation. The dilution series for each sample was done once and each dilution was plated in triplicate. The averages of colony counts expressed as colony forming units (cfu) per mL were calculated and plotted. Mating percentage was determined by dividing the cfu/mL from CA +ura agar by those from CA +ura/trp/thy agar. Samples of the H53 and H98 isolates used for the mating experiments were also plated on CA + ura as negative controls.

RNA was isolated from cells in Trizol reagent as per manufacturer’s instructions. The RNA from two filters was pooled for each sample. Isolated RNA was subjected to two rounds of DNAse digestion using Turbo DNA-free (Thermo Fisher) as per manufacturer’s directions. DNAse-treated RNA was finally purified using Qiagen RNeasy columns, as per manufacturer’s directions. RNA was assessed for DNA contamination using PCR, and RNA quality was determined using a 2100 Bioanalyzer (Agilent genomics). cDNA libraries for sequencing were constructed using a ScriptSeq Complete v2 RNA-Seq Library Preparation kit (Illumina), including depletion of rRNA using the Ribo-Zero rRNA removal beads for bacteria (a common practise for archaeal RNA sequencing ^27, 28^) and barcoded with the ScriptSeq Index PCR Primers (Set 1). cDNA was sequenced using a 150-cycle MiSeq reagent kit v3. Libraries were pooled and sequenced at 10 pM with a 15% 12 pM PhiX spike-in.

### Bioinformatics

All commands with specific parameters can be found at Gogarten-Lab github repository ^29^. Quality control on the sequences included Scythe v0.991 ^30^ adapter trimming against Illumina universal adapter sets followed by Sickle v1.33 ^31^ read trimming. Trimmed reads were aligned to the *Haloferax volcanii* DS2 genome (NCBI accession GCF_000025685.1_ASM2568v1; 4.0129 Mb length; 65.46 %GC; 4,023 genes) and quantified using Salmon v0.9.1 ^32^ with Variational Bayesian Estimation Method (VBEM). The *Haloferax volcanii* DS2 genome annotations were cross referenced against the UniProt database ^33^ and a complete table of the annotations and abundance estimates is available (Supplementary Table S3). Additionally, alignments were completed using bowtie2 v2.3.1 ^34-36^. Counts were normalized using trimmed mean of M-values (TMM) ^37^ and differential expression analysis was completed using NOISeq v2.26.1 ^38, 39^ in R by running noiseqbio function with default parameters. Differential expression analysis was completed for all comparisons ^29^ and comparisons between planktonic state and each time point are presented here. Interactive volcano plots and Manhattan plots were created in R using manhattanly v0.2.0. ^40^. Pearson and Spearman correlation coefficients were calculated using the cor function in R stats package v3.5.2. Principle component analysis was completed in NOISeq (Euclidian distance) and using r stats prcomp function with linear regression models calculated for PC1, PC2, and PC3. Bar charts and violin plots were constructed using ggplot2 v3.1.0 ^41-43^. Phylogeny for HVO_RS15650 (conjugal transfer protein) was constructed via NCBI; briefly, protein sequence was BLASTed against the non-redundant database, top 100 hits were BLAST pairwise aligned, using Fast Minimum Evolution tree method, and Grishin distance with maximum sequence distance of 0.85 ^44-46^. Genes were clustered using normalized expression counts via DESeq2 v1.24.0 ^47^ with a likelihood ratio test and an adjusted p-value cutoff of 0.05. Differential expression using NOISeq v2.26.1 was completed for cluster three and volcano plots were produced with Manhattanly v0.2.0. A homology analysis of the hypothetical proteins was conducted using HHblits v3.3.0 from the HHSuite3 package against the Uniclust30 database ^48, 49^. N-glycosylation residues were predicted using the NetNGlyc 1.0 server ^50^. Additional data preparations and organizing were completed in R using tidyr ^51^, dplyr ^52^, and reshape2 ^53^. The graphical outline in figure 1 was constructed using biorender.com. Reads were deposited to NCBI under BioProject PRJNA632928 and SRA accession numbers can be found in Supplementary Table S2.

## 3. Results and Discussion

### 3.1 Mating and mating efficiency

Cells were mated by mixing the H53 and H98 auxotrophs in equal volumes and transferring the cells onto filters. H53 cells are auxotrophic for tryptophan and H98 cells are auxotrophic for thymidine. The genes *trpA* and *hdrB* ^26^ were thus used as markers for mating. Cells able to grow on CA plates that were supplemented with neither tryptophan nor thymidine represent mated cells. The number of colony-forming units (cfus) on plates supplemented with uracil divided by cfus on plates supplemented with uracil, thymidine, and tryptophan yielded a measure of mating efficiency at each time point ^26^. We note that this measure excludes the majority of mating events: all mating between two H53 and between two H98 go undetected and mating between H53 and H98 are only detected if the heteroploid state is resolved after a recombination event in a way that results in a hybrid chromosome with both functional genes. Nevertheless, this measure is a standard means of gauging mating efficiency ^15, 18, 19, 54^. Mating for planktonic cells over time is not shown because under our experimental conditions we saw no mating in planktonic cells. Even though mating in planktonic cells has once been reported, rates were 10^4^ times less than for cells on filters, so mating in planktonic cells can be considered negligible under our experimental conditions ^55^. Samples from the original H53 and H98 cultures used for the mating were also plated separately on CA + ura as a negative control and no growth was observed on these plates. That, coupled with no observed mating in the planktonic cells, indicates a negligible rate of false positives.

Mating was allowed to proceed by incubating the filters on solid medium and cells were harvested for RNA isolation and plating at various time points (see Materials and Methods and Fig. 1A). The intention of this paper was to observe changes in gene expression between non-mated and mated cells over time, but likewise some changes might also reflect the cells’ reaction to moving from a suspended to sessile milieu. Additionally, the total cfu count did increase over time, suggesting the cells were dividing (Fig. 1B) with a doubling time below 4 hours. While this is in agreement with some sources ^56^, others consider this rapid ^57^. Given the rather loose adherence of the typical *Hfx. volcanii* biofilm ^16, 58^, the increase in cells could be representative of a combination of cell growth as well as more of the cells sloughing off filters at later time points. Regardless, some gene expression level changes could be in response to growth and changes in local nutrient availability.

Transcriptome data were taken from cells exposed to conditions that encourage mating, but there was no means of specifically selecting out mating cells from this group. Nonetheless, we feel that this data is a means of surveying for genes that may have a significant role in the cellular processes surrounding mating. Mating was observed to start rapidly, as even those cells harvested immediately were shown to produce colonies able to grow on unsupplemented CA medium (Supplementary Table S1). It is unlikely that mating is completed so rapidly, but the mating at 0 hours could be indicative of the onset of mating. In other words, the cells were able to adhere to one-another to initiate mating immediately, and this adherence was robust enough that the mating was able to continue to completion as the cells were washed from the filters where mating was initiated. Over the course of 24 hours, mating efficiency rose to approximately 4 x 10^−5^ comparable to that of previous work ^15, 19^. The main increase in cfus and mating efficiency occurred during the first 2 hours. While the mating efficiency for both replicates is in good agreement for most time points, those for 4 hours of mating are divergent, increasing to 7.8 x 10^−5^. Principle component analysis of the expression data (Fig. 1D), however, shows excellent agreement between the replicates in all time points.

### Gene expression overview

The transcripts were aligned to 3,941 genes in the *Hfx. volcanii* genome and expression was identified in 3,582 genes after removing genes with a total expression estimate of less than ten transcripts. Significant differential expression (p-value < 0.01) for each time point against the planktonic state transcriptome ranged from 109 genes in the zero-hour time point up to 1,092 genes at the four-hour time point (Table 1). Many of those genes are related to cell growth and moving from a planktonic to sessile state, while others may be associated with processes directly or indirectly related to mating. Genes with the highest and lowest log2 fold change were identified and dominated by hypothetical proteins (Supplementary Fig. S7, Supplementary Note). Those genes listed as hypothetical proteins were further analyzed for homology to genes of known function and described (Supplementary Note). Mating occurs through stages including cell contact, cell surface fusion, a heteroploid state, and separation into hybrid cells ^59^. There are genes which may contribute to or limit mating at each of these steps, so the focus was on the genes, pathways, and systems hypothesized to be involved in each of these processes and on genes that show a similar expression pattern to those involved in mating.

**Table 1.** Sample information including experimental parameters, sequencing and assembly information

| Sample ID | Treatment<sup>a</sup> | Raw Pairs<sup>b</sup> | Trimmed Pairs<sup>c</sup> | %remaining | Bowtie2<br>alignment rate<sup>d</sup> | No. D.E. genes<sup>e</sup> |
| --- | --- | --- | --- | --- | --- | --- |
| shaking | planktonic | 3,169,158 | 2,918,605 | 92.09% | 91.50% | n/a |
| t00 | 0 hours | 3,256,394 | 2,805,158 | 86.14% | 91.61% | 3,320 (109) |
| t02 | 2 hours | 3,036,028 | 2,789,144 | 91.87% | 94.17% | 3,434 (941) |
| t04 | 4 hours | 3,155,477 | 2,900,100 | 91.91% | 93.50% | 3,480 (1092) |
| t08 | 8 hours | 3,054,785 | 2,756,910 | 90.25% | 94.08% | 3,472 (757) |
| t24 | 24 hours | 3,136,767 | 2,873,436 | 91.61% | 93.27% | 3,462 (643) |

<sup>a</sup>Shaking represents cells suspended and not mating; each hour following is time on filter before sampling (see methods for details)
<sup>b</sup>Paired end sequencing (2x75) using Illumina MiSeq summed for two replicates
<sup>c</sup>Trimmed using Sickle v1.33 summed for two replicates
<sup>d</sup>Trimmed reads were aligned back to the *Haloferax volcanii* DS2 genome (NCBI assembly: ASM2568v1); average across two replicates
<sup>e</sup>Total number of differentially expressed genes for each time point compared to shaking control and genes with p-value < 0.01 in parenthesis

### Glycosylation and low salt glycosylation genes

While little is known about which genes are involved with mating of *Haloferax* spp. cells, it was previously noted that mating efficiencies were greater within than between species, though inter-species mating still occurs and the barriers to this are relatively lax. More recent work was done to determine if recognition of potential mating partners might be mediated in part by the glycosylation of cell-surface glycoproteins. Briefly, it was shown that disruption of normal glycosylation by either gene knock-out or culturing cells in low-salt conditions can reduce mating efficiencies ^19^. It is for this reason that glycosylation and low-salt glycosylation gene expression levels were specifically examined. In addition to those known genes in the N-glycosylation pathways, a homologous gene analysis of the hypothetical proteins was completed to identify potential additional genes involved in N-glycosylation (Supplementary Note) and a search of predicted glycosylated residues was conducted and made available on github ^29^.

Overall, for the majority of both glycosylation and low-salt glycosylation genes, expression increased when compared to suspended cells (Fig. 2). Most of the N-glycosylation and low salt N-glycosylation genes showed an expression pattern reflecting time of mating – increased expression at 2-4 hours then gradually decreasing – with consistent and significant observations (also see section regarding the identification of additional mating associated genes via gene expression patterns, below). The two-hour time point had significantly higher expression in eleven genes involved with glycosylation with the highest expression for the major cell surface glycoprotein, HVO_RS14660 (fold change > 16), generally one of the highest expressed genes in *Hfx. volcanii* DS2. Of the glycosylation genes that were previously investigated in relation to mating ^19^, *aglD* (HVO_RS08530) shows an increase in expression beginning at 2h of mating, and *aglB* (HVO_RS12050) shows a small increase in expression levels at 0h with a substantial and significant increase beginning at 2h of mating, although in the former case only the changes in gene expression at 2h of mating are statistically significant (p<0.05). The low-salt glycosylation gene *agl15* (HVO_RS20240) showed a significant increase in expression levels in comparison to that of planktonic cells in all time points after 0 hours of mating (p<0.05).

**Figure 2:**
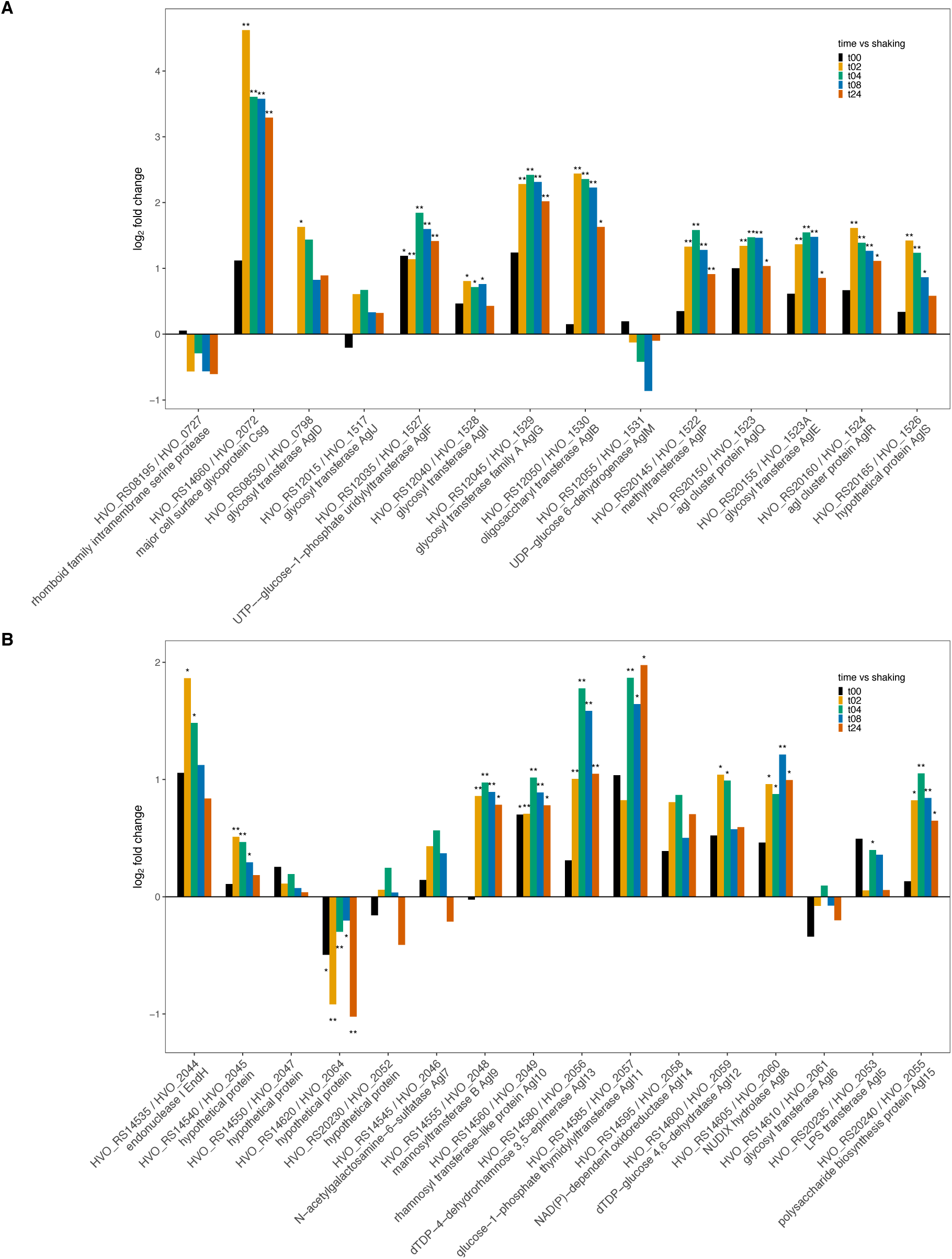
Bar plot depiction of differential expression compared to planktonic state for (A) glycosylation genes and (B) low-salt glycosylation genes; gene id on the x-axis and log2 fold change on the y-axis, *p-value <0.05 and **p-value<0.01. Genes directly involved in the N-glycosylation pathways are designated with their “*agl*” gene name and the remaining genes have a putative connection to glycosylation.

In the work done by Shalev et al., the deletion of both the *aglB* and *agl15* genes showed a large negative impact on mating efficiency, with deletion of both genes together reducing mating to almost undetectable levels ^19^. The expression data agrees with this observation, showing an immediate increase and tapering over time. Deletion of the *aglD* gene had a very small impact on mating efficiency with even a small increase shown when both mating partners had the gene deleted ^19^. Both *aglB* and *agl15* are highly significant, while *aglD* is only statistically significant at the 2h time point. The pattern is somewhat similar to *aglB*, but not to *agl15*, which peaks at 4h, not 2h. Other genes known to have an impact on cell-surface glycosylation (e.g., *aglF, aglG*, and *aglI*; HVO_RS12035, HVO_RS12045, and HVO_RS12040, respectively), but with as-yet undescribed impact on mating, show significant increases in expression in mated cells compared with unmated cells.

The levels of expression of many of the genes in the low salt glycosylation pathway increased even though the mating was performed in salinities above “optimal” in the mating studies by Shalev et al., and well above the concentrations used in the “low salt” conditions ^19^. While, expression of low salt glycosylation machinery under conditions of normal salinity have been previously observed ^60^, in this case the initiation of the pathway could be in response to perturbation of the cell membrane, whether caused by decreased salinity or other potential stresses, such as viral infection ^21^. One explanation for our transcript data would be that the formation of cytoplasmic bridges connecting cell envelopes during mating may be an additional membrane stress factor affecting “low salt” glycosylation gene synthesis.

These data overall suggest that while expression levels already existing in cells when they are subjected to mating conditions may have an impact, the mating itself may also have an impact on the expression of these genes. Conversely, it may not be necessary for the cells to increase the expression of genes related to glycosylation to initiate mating. The changes observed may therefore represent fluctuations resulting from the cells’ growth or, most likely, a change from planktonic to sessile cells. The observation that genes whose deletions have a positive impact on mating and those who have a negative impact on mating both show increases in expression post-mating implies that increased glycosylation is a physiological response to growth on a surface that probably affects multiple processes including mating.

### Community Development and Biofilm Formation

*Hfx. volcanii* is known to form biofilms ^16, 58^ and incubating cells after filtration will form a biofilm over time. A lack of observable mating in planktonic cells (Supplementary Table S1), combined with prior observation of an increase in mating in cells which are part of a biofilm ^16^, suggests that the majority of mating in the natural environment will take place amongst *Hfx. volcanii* that are part of a biofilm. This may be mediated in part by molecules involved in cell adhesion, for example the glycosylation genes already discussed, as well as biofilm formation and putative quorum sensing genes. As such, we examined which transcripts predicted to be involved in those processes increased or decreased in comparison to planktonic cells (Fig. 3).

**Figure 3.**
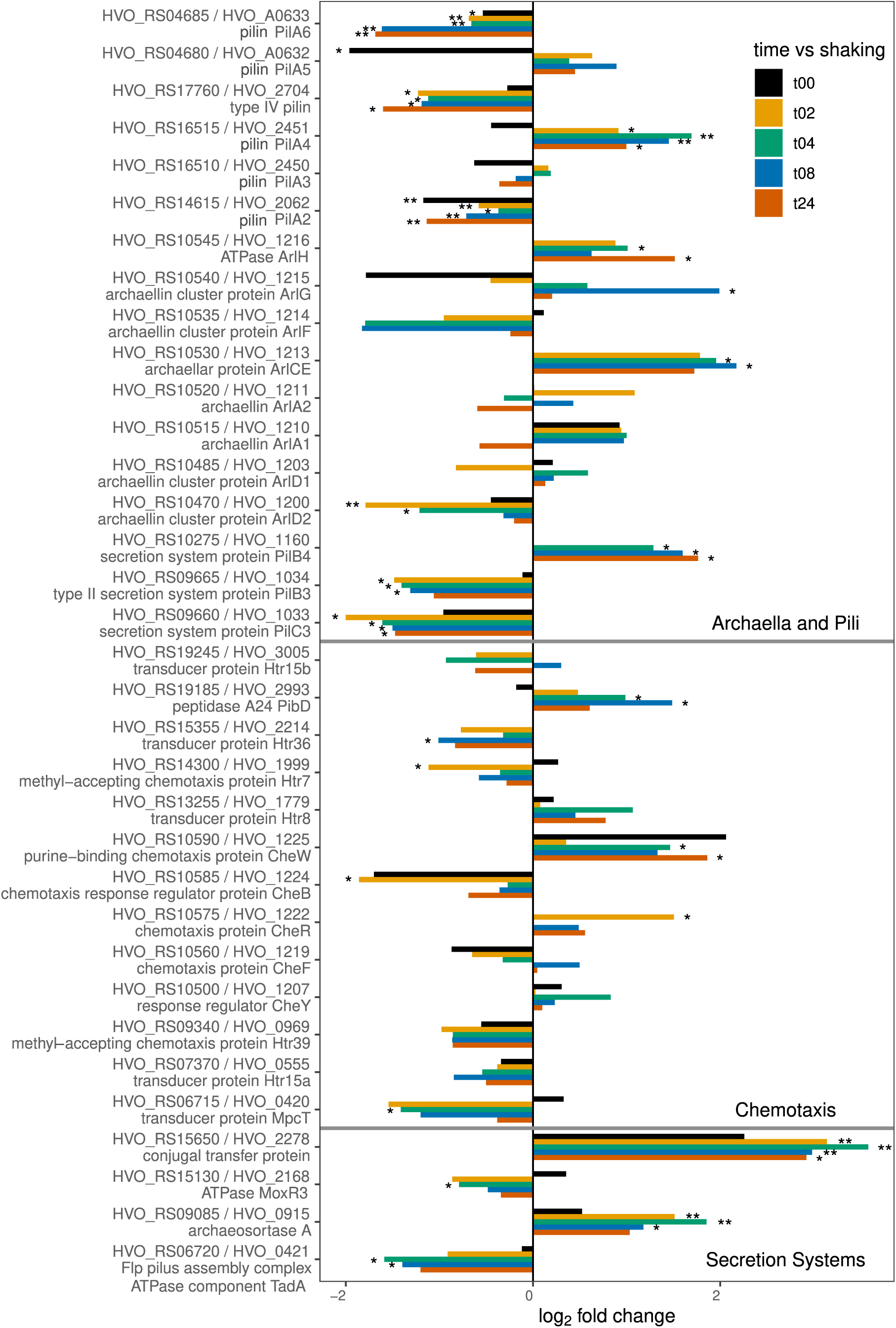
Bar plot depiction of differential expression compared to planktonic state for genes involved in community development and biofilm formation; gene id on the y-axis and log2 fold change on the x-axis, *p-value <0.05 and **p-value<0.01.

Biofilm formation begins with adhesion of cells to a surface. Archaeal flagella, also referred to as archaella ^61, 62^, although sharing a role comparable to that of bacterial flagella, are assembled in a manner more similar to bacterial type IV pili ^63-65^ and some proteins involved in the processing of both pilin and archaellins are shared ^55, 66^. Archaella and chemotaxis proteins modulate motility of cells to find alternative conditions in their aqueous environment and allow cells to move to surfaces ^55, 58^. It has been suggested that in the case of bacteria, flagella may play an important role at the onset of biofilm formation ^67^. While it has been shown that the *Hfx. volcanii* archaella are not important for adhesion ^55^, the regulation of both archaella and archaeal pili may affect biofilm formation and mating and their chemotaxis proteins have been previously implicated in mediating social mobility within a biofilm ^16^. Archaella and pilin assembly proteins had non-monotonous transcriptional changes (Fig. 3). The biogenesis components, *pilC*3 and *pilB*3 (HVO_RS09660, and HVO_RS09665, respectively) ^66, 68^, had significantly lower expression with trending decreases in fold-change (Fig. 3). The use of the archaella for movement in the planktonic condition is supported by the decreased expression over time. However, some of the genes associated with motility and the regulation of motility ^69, 70^, including *arlCE, pilA4, pilA5* (HVO_RS10530, HVO_RS04680, and HVO_RS16515, respectively), showed increased expression with some statistical significance.

While there is no notable expression pattern across all chemotaxis genes, many time points do show significant increases and decreases in expression. For example, *cheW* (HVO_RS10590), coding a chemotaxis adaptor protein that shares a role in stimulating the rotation of the archaella ^71^, has a significant increase at 4 and 8-hour time points, while *cheB* (HVO_RS10585), a methylesterase that modulates chemotaxis by affecting archaellar rotation ^72, 73^, is significantly decreased at 2 hours and moderately decreased at all other timepoints (Fig. 3). Genes used for motility specifically would be expected to decrease in expression once cells have adhered to a surface, while those used for adherence would be expected to increase temporarily until the biofilm structure is formed. This speculation is supported by observations in the formation of bacterial biofilms ^67^. When *Hfx. volcanii* archaellin genes *arlA1* and *arlA2* (HVO_RS10515 and HVO_RS10520, respectively) and the prepilin peptidase gene *pibD* (HVO_RS19185) were examined for their roles, deletion of the archaellin genes showed no impact on surface adhesion or on mating efficiency, but deletion of *pibD* did reduce adhesion to surfaces and, again, mating efficiency was unaffected ^55^. Neither of *arlA1* and *arlA2* had any significant changes in transcription levels, but *pibD* gene transcription levels increase in response to transfer to mating conditions with statistically significant increases at 4 and 8 hours (Fig. 3). This does not contradict previous studies showing *pibD* does not to have an impact on mating and the increase may in fact reflect the role of *pibD* in processing type IV pilins ^55, 66^.

Biofilm adhesion is also facilitated by the presence of extracellular polymeric substances (EPSs) made up of DNA, amyloids, and glycoproteins, which together make up the biofilm structure ^58, 74-76^. EPS is produced by the cell and transported utilizing secretion system machinery. Archaeosortase genes, for example, mediate the attachment of proteins to the cell surface and have been shown to impact not only the cell growth and motility but also the amount of mating between cells ^18^. Deletion of the *artA* archaeosortase gene (HVO_RS09085) decreases mating ^18^, an observation that correlates well with the transcriptome data: *artA* increases at all timepoints with peak expression at 4 hours and statistical significance from 2-8 hours (Fig. 3). HVO_RS15650, which has been annotated as both a “type IV secretion system DNA-binding domain-containing protein (TraD)” and as a “conjugal transfer protein”, had highly significant increases in expression, also with peak expression at 4 hours (Fig. 3, Supplementary Fig. S1). In bacteria, type IV secretion systems (T4SS) are responsible for conjugation in some species, as well as a variety of other roles, including delivery of effector molecules to eukaryotic hosts, biofilm formation, and killing other bacteria ^77^. The phylogeny of HVO_RS15650 and its homologs shows those belonging to archaeal species are in a clade separate from the bacterial homologs (Supplementary Fig. S2), thus one cannot assume a conjugation-related role for this protein with certainty. While it will be interesting to see if this gene has a further role in the mating process in Archaea, the observed expression pattern could be due to a role in biofilm formation. Many putative secretion system genes appear to have lower expression levels in the filter-bound cells and many expression patterns are variable with rapid increases and decreases consistent with biofilm maturation (Fig. 3, Supplementary Fig. S1). Overall, archaeosortase and type IV secretion system genes are consistent with our hypothesis for expression changes associated with a mating environment.

### DedA/SNARE genes

Fusion processes have long been studied in eukaryotic systems in intracellular trafficking and vesicle maturation, viral entry, and gamete fusion ^78-80^. Similar mechanisms appear to be at play in *Hfx. volcanii* mating which involves fusion and cleavage of membranes of neighbouring cells ^14, 15, 59^. SNARE proteins have been implicated in eukaryotic vesicle fusion in mammalian neuronal cells and similar machinery and processes have also been described in yeast ^81^. The SNARE-associated Tvp38 family of proteins in yeast have homologs in other single- and multi-cellular eukaryotes and are homologous to the DedA protein family in bacteria and archaea ^82-84^. While the Tvp38 proteins do not appear essential in yeast, the DedA family of proteins in bacteria has been implicated in division defects causing long chains of incomplete cells to form ^85^. While the proteins implicated in cell division are initially appropriately localized at a site of division, they disassemble again when division fails ^85^. Of archaeal genomes examined, 27% lack genes with significant sequence similarity to DedA, but all Halobacteriales examined contain a DedA homolog ^79^. Given the proposed manner of cell mating (partial or whole cell fusion followed by cleavage), it is reasonable to predict that DedA- or SNARE-type proteins will have a role in this process.

Three DedA family proteins found in *Hfx. volcanii* show appreciable changes in transcription (Fig. 4). HVO_RS01005 shows an immediate increase in transcription at the 0-hour time point, lower increases at 2 and 4 hours, and higher increases at 8 and 24 hours of mating, but none of the differential expression is statistically significant (p > 0.05). HVO_RS05125, a hypothetical protein with a DedA motif, also shows an immediate increase in transcription at the zero time point, but with decrease transcription thereafter. Only the decreases at 2 and 24 hours are significant (p > 0.05). Another DedA homologue, HVO_RS09165, shows an immediate drop in expression (p = 0.059) followed by significantly lower expression at 2, 4, 8, and 24 hours (p-values: 0.003, 0.006, 0.010, and 0.005 respectively). HVO_RS09165 is considered to be a member of the TVP family of proteins, which are yeast SNARE proteins conserved in numerous eukaryotes ^84^. It is intriguing that the expression of the TVP family member protein expression is in the opposite direction expected for genes involved in mating, and it could indicate that the transient fusion of cells during mating is mediated by the balance of DedA proteins present before cells are transferred to mating conditions.

**Figure 4.**
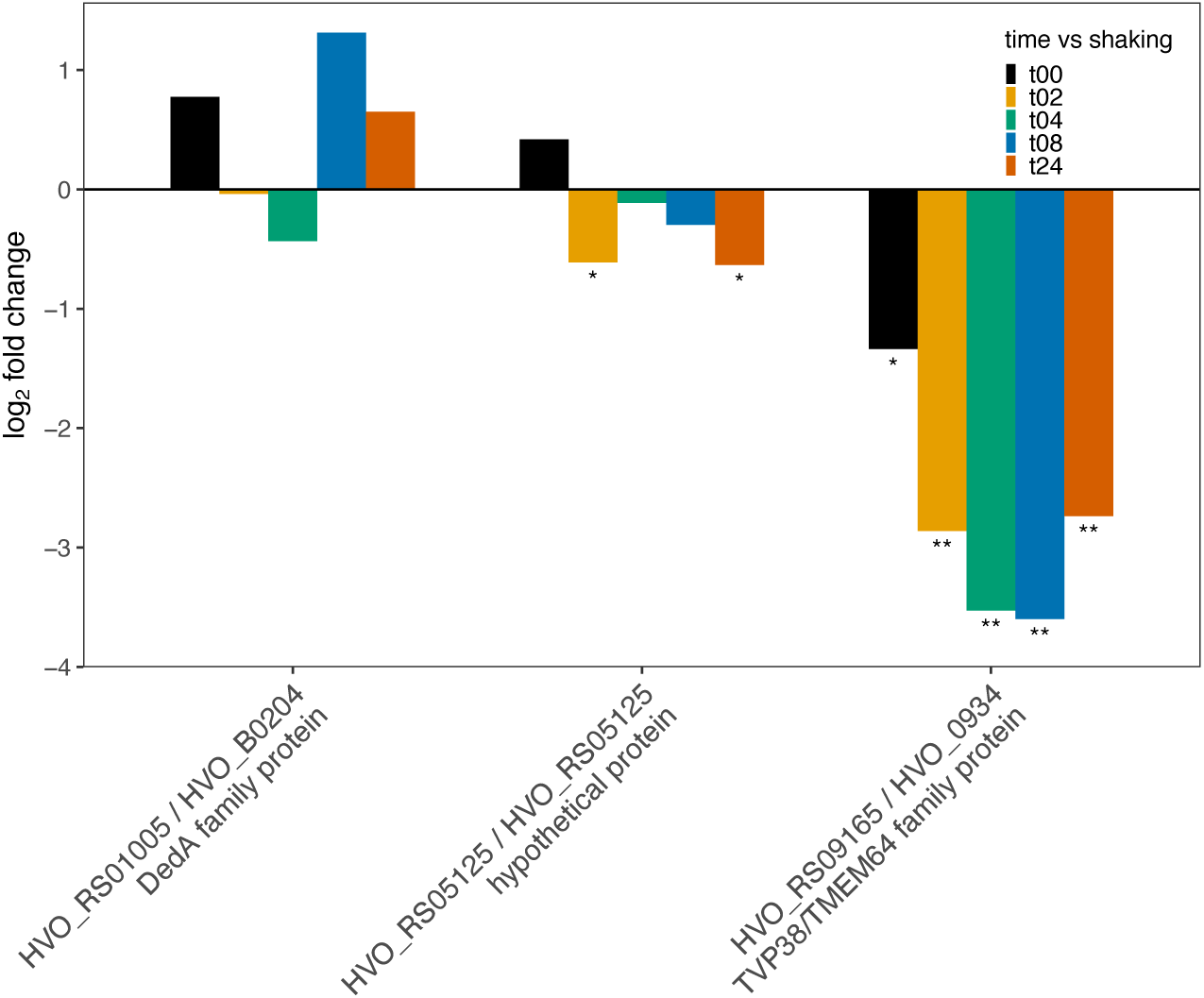
Bar plot depiction of differential expression compared to planktonic state for SNARE/*dedA* motif genes; gene id on the x-axis and log2 fold change on the y-axis, *p-value <0.05 and **p-value<0.01.

### CetZ/FtsZ

As can be seen in Fig. 1A, cells that have been incubated on filters for mating also undergo cell division over the course of 24 hours. It is probable that there is an overlap in machinery used for cell division and mating, in particular, those genes responsible for DNA replication and the cytoskeletal proteins. In mating, *Haloferax* are known to exchange large segments of both genomic and plasmid DNA ^15^ and passive diffusion of DNA through a cell is limited by size ^86^, so one can postulate that the movement of such large molecules requires the involvement of the cytoskeletal matrix. Research in eukaryotes has shown the requirement of the cytoskeleton for trafficking of DNA to the nucleus in transfected cells ^87^ and it is conceivable that an analogous system may be at play in the transport of DNA between mating *Haloferax* cells. Additionally, the CetZ proteins appear to mediate the shape of *Haloferax* cells ^88^, and thus might be responsible for the formation of cytoplasmic bridges. For these reasons, the expression levels of the *cetZ* and *ftsZ* genes were investigated.

CetZ1 has been implicated in cell motility in *Hfx. volcanii* ^88^. Surprisingly, transcriptome data showed that expression of CetZ1 (HVO_RS15305) was increased in sessile cells as compared with planktonic cells (Fig. 5). While Duggin et al. ^88^ showed that CetZ1 converted cells to the rod-shape required for motility, the observed increase in expression may indicate that the protein is also required for mating. Indeed, many of the *cetZ* genes show increased expression levels under mating conditions with the exception of *cetZ2* (HVO_RS08275) and *cetZ3* (HVO_RS10050). Both *cetZ2* and *cetZ3* show decreased expression at most time points, the exception being at 0-hour wherein *cetZ2* shows an increase in expression and *cetZ3* shows a decrease in expression. Both these genes are tubulin-like genes and may have a role in cell shape or rigidity ^88^, but their exact function is not yet clear. While it is tempting to speculate that both the rigidity of the cell shape may play a role in mating and its regulation by the increase and decrease of the expression of different *cetZ* genes could be important, further understanding as to the function of these proteins is essential.

**Figure 5:**
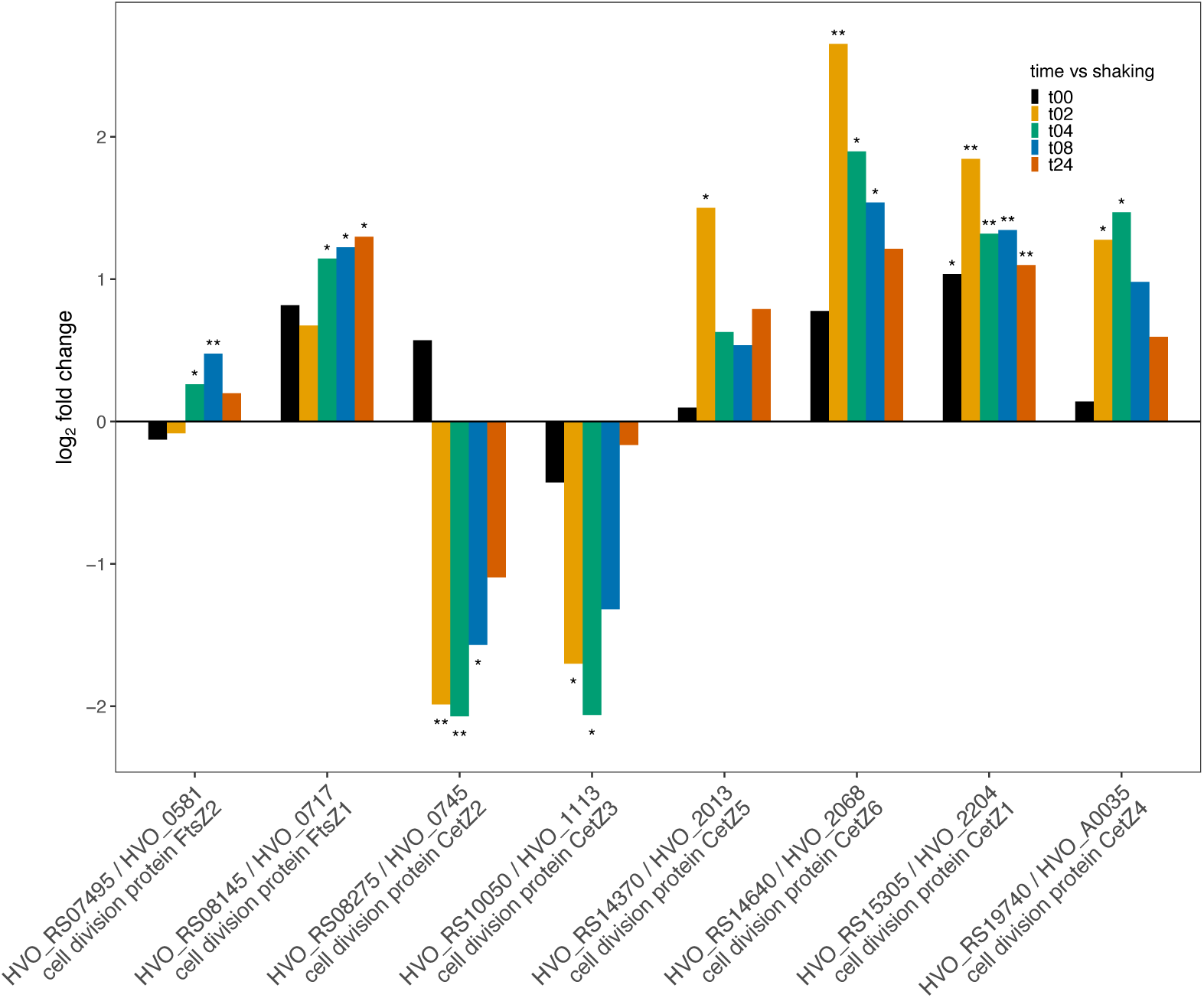
Bar plot depiction of differential expression compared to planktonic state for FitZ and CetZ genes; gene id on the x-axis and log2 fold change on the y-axis, *p-value <0.05 and **p-value<0.01.

In addition to the *cetZ* genes, *Hfx. volcanii* also has the genes *ftsZ1* and *ftsZ2* (HVO_RS08145 and HVO_RS07495, respectively). These genes have been implicated in cell division: the encoded proteins form a ring structure that effectively pinches off dividing cells from one-another ^89, 90^. The *ftsZ1* gene is also well-conserved amongst many Haloarchaea ^91-93^ and both *ftsZ1* and *ftsZ2* have recently been shown to be essential for proper cell division in *Hfx. volcanii* ^94^. In the case of this mating transcriptome, *ftsZ1* shows higher rates of transcription in the mated cells as opposed to the unmated, but *ftsZ2* transcription rates are slightly lower at 0- and 2-hour time points in mating and have small increases in later time points that are significant at 4 and 8 hours (Fig. 5). While it is conceivable that the same mechanism of separation of dividing cells is used in the separation of mating cells, the growth and division of cells after plating may also account for the increase in transcription of *ftsZ1*. As yet, understanding of the roles of the *cetZ* and *ftsZ* proteins in *Haloferax* is limited and further research into their roles in mating and cell functions are warranted.

### Selfish Genes

*Hfx. volcanii* is known to harbor selfish genes, as well as innate and adaptive means of defense against their invasion ^95, 96^. Studies in natural populations have shown a larger amount of HGT within than between haloarchaeal phylogroups ^97, 98^, which could, to some degree, be attributable to barriers to mating which are active after cell fusion. Although the *Hfx. volcanii* cells were mated to members of the same species in this experiment, it is possible that the act of mating might trigger expression of selfish genes and those associated with defense against invasion of selfish genetic elements.

The *Hfx. volcanii* transcriptome contains expressed transposases from at least nine insertion sequence (IS) families on the main chromosome and on plasmids pHV1, pHV3, and pHV4 (Fig. 6A). Transposase genes can have high similarity to one another. To ensure that the expression output was as accurate as possible, the alignment program, Salmon, was chosen. Salmon initially removes exact duplicates at indexing of the reference genome, then uses a dual-phased statistical approach to robustly estimate alignment scores. An overwhelming majority of IS families show increased expression at all or most time points compared to the shaking control; forty-two have significantly higher expression, while only ten have significantly lower expression (Fig. 6A, Supplementary Fig. S3, Supplementary Table S5), supporting the hypothesis that selfish genes entering a mating condition will be activated. The IS5 and IS6 families’ expression, for example, are almost entirely increased after mating (Supplementary Fig. S3, Supplementary Table S5). While most of the insertion sequences have higher expression than the shaking control, there are still many expressed during shaking and a few with expression decreased in a sessile state (Fig. 6B, Supplementary Fig. S3, Supplementary Table S5). One family, the IS110 transposases, all have higher expression during shaking (Supplementary Fig. S3, Supplementary Table S5). The largest family of transposases (IS4) has both increased and decreased fold changes, but is mostly expressed after plating, even at the zero-time point.

**Figure 6:**
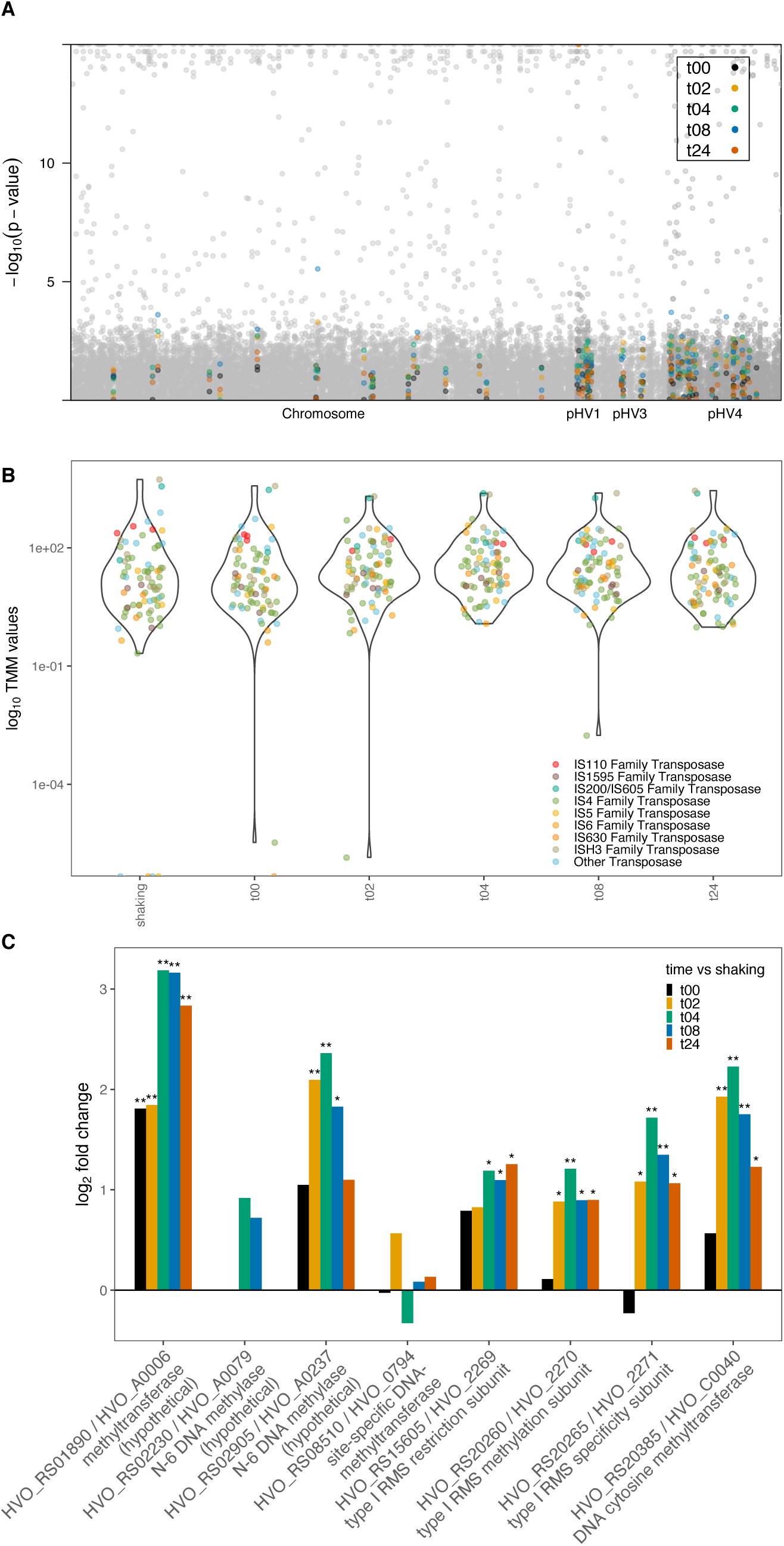
(A) Manhattan plot with chromosome number on the x-axis and -log10 p-value of differential expression for planktonic cells vs plated timepoints on the y-axis, all genes are plotted in grey and genes for insertion sequences are highlighted by color according to the key; (B) Violin plot showing log_10_ of average normalized expression values for each treatment on the y-axis; (C) Bar plot depiction of differential expression compared to planktonic state for restriction modification system genes; gene id on the x-axis and log2 fold change on the y-axis, *p-value <0.05 and **p-value<0.01.

Although it has been previously observed that the expression of CRISPR-associated genes is low in *Haloferax* species under normal growth conditions ^99^, it has been shown that not only are CRISPR spacers acquired during cross-species mating, but that such spacer acquisition will reduce the gene transfer during cross-species mating ^54^. All the CRISPR-associated genes in *Hfx. volcanii* are located on pHV4 (Supplementary Fig. S4A) ^100^. All *cas* genes were expressed at all time points (Supplementary Fig. S4B), but changes in expression levels varied. Two genes that encode components of the interference machinery, *cas8b* and *cas7* (HVO_RS02760 and HVO_RS02765, respectively), had significantly increased expression from 4-8 hours after plating. Two genes, *cas6* and *cas2* (HVO_RS02755 and HVO_RS2790, respectively), the former required for processing of CRISPR RNAs, and the latter involved in acquiring new spacers, were expressed significantly less than in planktonic cells starting at the 2-hour time point to 24-hours and 8-hours, respectively. *cas4* (HVO_RS02780), also assumed to be a part of the spacer acquisition module, had a significant increase in expression immediately at 0-hour turning to lower expression for the rest of the experiment. Acquisitions of spacers were shown to be induced by mating with different *Hfx*. species, e.g. *Hfx mediterranei*, but spacer acquisition also occurred during within-species mating, albeit to a lesser extent ^54^. It therefore appears that while CRISPR-surveillance against threats (i.e. interference module) is maintained or even increased during mating conditions, spacer acquisition remains generally low, potentially preventing acquisition of DNA spacers from closely related cells during mating, which could result in CRISPR autoimmunity.

Another potential limiting factor on genetic exchange in *Haloferax* during mating is the activity of restriction modification (RM) system enzymes, provided the mating pair possess different sets of RM systems. RM systems also constitute addiction cassettes, and therefore can be considered selfish genetic elements (see Fullmer et al. ^101^ for discussion). *Hfx. volcanii* contains a Type I RM system and a number of DNA and RNA methylases ^101-103^. Strains of *Hfx. volcanii* auxotophs H53 and H98 were constructed which lack most of the original RM system genes previously described ^102, 103^. Studies by Ouellette et al., (2020) ^104^ suggest that frequency of recombination during mating was lower for strains with incompatible RM systems than those with compatible RM systems, although the effect could have been accounted for by the presence of an RM motif close to one of the marker genes ^104^. In contrast, mating a strain harboring an intein with a homing endonuclease and a strain without the intein, resulted in increased recombination rates ^105^, revealing that DNA double strand breaks can lead to increased recombination during mating. All RM genes show increased transcription during mating conditions in comparison to planktonic cells with the only exception being an initial decrease at the 0-hour time point for the gene in the *rmeRMS* operon (consisting of the genes HVO_RS15605, HVO_RS20260, and HVO_RS20265) encoding the specificity unit (HVO_RS20265) (Fig. 6C). The transcription of the genes HVO_RS01890 and HVO_RS02905, predicted methyltransferases, are increased in response to mating, although HVO_RS02905 sees an increase to a lesser degree than that of HVO_RS01890. While it has been suggested that these two genes were possibly at one point a single RM system ^102^, a later study showed that neither of them are in fact functional in *Hfx. volcanii* ^103^. The same study showed that all methylation within *Hfx. volcanii* is mediated by the gene HVO_RS08510 and the operon *rmeRMS*. HVO_RS08510 is an orphan methyltransferase and is responsible for the methylation of a CTAG motif ^103^, which is widely distributed amongst the Haloarchaea ^101^. This gene shows some inconsistent variation of transcription in mating cells as compared to planktonic cells, but none of the changes observed are significant and expression values are low (Supplementary Table S3). The *rmeRMS* operon was shown to be responsible for all the adenine methylation in *Hfx. volcanii* ^103^, and it encodes a type I restriction-methylation system. The transcription of all three genes showed an increase after the 0-hour time point of mating. The genes HVO_RS02230 and HVO_RS20385, were identified as methyltransferase homologs, but were not functionally active ^103^. Both these genes also showed an increase in transcription in mated cells as compared to planktonic cells, although in the case of HVO_RS02230, this increase only appeared in the 4- and 8-hour time points. Increased expression of RM system genes might be selected for because it increases vigilance against selfish genetic elements or because an increased restriction endonuclease activity guards against deletion of the RM system genes through recombination with DNA from a strain that does not harbor the particular RM system. Overall, these results coupled with the study showing increased mating between *Haloferax* cells with compatible RM systems suggests that RM systems may play a role in limiting recombination after cell fusion.

### Identification of additional mating associated genes via gene expression patterns

Beyond our hypothesized or known genes for mating, we were interested in identifying additional unknown genes that could also play a role in mating. To that end we applied a clustering analysis to group genes based on expression changes across the time points resulting in clusters of genes with shared expression patterns. Clustering analysis based on expression patterns resulted in twenty-two clusters with a range of 3 to 289 genes per cluster (Supplementary Fig. S5). Those genes that increased from shaking or time 0 rapidly to the two-hour time-point and then decreased in expression after two hours show a pattern consistent with the timeline for mating (Supplementary Fig. S5; Supplementary Table S4). Clusters with continuous increases to twenty-four hours, such as the pattern seen in clusters 2 and 16, are less likely to be associated with mating and more likely associated with cell growth on solid medium. This is because mating is initiated predominately within the first two hours of plating (Fig. 1B,C). Cluster three contained the most genes of all clusters and also presented the putative mating pattern of gene expression.

Differential expression analysis on cluster three between the shaking control and time points 2-24 hours revealed a number of significant genes, many of which were also found in the top 10 differentially expressed genes for the 2, 4, and 8-hour time points (Supplementary Fig. S5, Supplementary Fig. S6). Most of the highly significant genes in cluster three in the differential expression analysis comparing shaking or t0 with the two- and four-hour time-points were ribosomal proteins or hypothetical proteins (Supplementary Table S4). Cluster three also contained a number of potential operon genes. Notably, co-located genes linked to glycosylation were identified – HVO_RS20145, HVO_RS20160, and HVO_RS20165 (Supplementary Fig. S8). These genes are *aglP*, a SAM-dependent methyltransferase ^106^, *aglR*, which is involved in dolichol phosphate-mannose processing and possibly contributing to the flippase activity in the cell ^107^, and *aglS*, a dolichol phosphate-mannose mannosyltransferase ^108^, respectively. Cluster twelve, which has a similar expression pattern to cluster three but peaking at four-hours instead of two, contained additional genes in this location, HVO_RS20150 (*aglQ*, a glycosol hydrolase) and HVO_RS20155 (*aglI*, a glycosyl transferase). The genes share the expression profile described for a number of glycosylation genes (see previous section on glycosylation and Fig. 2).

This analysis suggests a large number of potential targets for further examination for their role in mating. The majority of glycosylation genes have not been experimentally examined for their role in *Hfx. volcanii* mating. There were also genes annotated as ABC transporter genes, multiple transposases, transcriptional regulators, and cell division proteins that could potentially be involved in various phases of mating. In addition to laboratory based knockout experiments for genes of known or presumed function, there is a gap in functional annotation that would benefit from additional computational examination.

## 4. Summary and conclusions

This work represents an overview of the gene expression changes between unmated *Hfx. volcanii* and cells grown under mating conditions. It has raised many avenues for further work into genes controlling mating behavior and likewise has suggested genes which may have a role in biofilm formation. One of the more remarkable observations is that the process of mating begins rapidly in *Hfx. volcanii*. While prior published mating experiments on filters include incubation times of 16 to 48 hours to allow for mating ^15, 18, 19, 54, 55^, here we show that cells initiate mating immediately after contact. Mating efficiency appears to peak between 4 to 8 hours after initiation, with very little difference between those two time points. Future experiments involving mating may be better streamlined in light of this.

Glycosylation has been previously examined for its connection to mating and is likely driving mating through S-layer decorating that serves to attract cells to one another. Disruption of glycosylation genes has resulted in reduction of mating ^19^, but no other genes have been distinctly linked to mating to date. Glycosylation genes were examined and followed a pattern consistent with the mating efficiency over time. Cells cultured in low-salt were also found to decrease mating efficiency ^19^ and the patterns of expression of many low-salt glycosylation genes were consistent with mating, though some were more erratically expressed. Multiple glycosylation genes were identified in this study for future examination on their impact on mating, including *aglF, aglG*, and *aglI*.

Glycosylation of the cell wall influences mating as the cells move closer together, a condition most often seen in biofilm formation and community development. Indeed, cells are observed to mate in biofilms at a significantly higher rate than in the planktonic state ^16^. Neither the genes involved in construction of the *Hfx. volcanii* archaella and pili, nor those involved in chemotaxis showed a uniform pattern of expression. Additionally, one previous study showed that disruption of the archaellin had no effect on mating efficiency ^55^ and genes associated with biofilm formation, namely adhesion, showed no effect on mating efficiency. Previously published gene deletion data in conjunction with our findings suggests that the genes associated with biofilm formation and community development are not driving mating, but instead probably generate the proximity necessary for mating in nature.

The physiology of mating in *Hfx. volcanii* involves cellular fusion via the formation of a cytoplasmic bridge followed by bidirectional transfer of DNA and ends with cell cleavage. SNARE/DedA family proteins in Haloarchaea do not have a clear transcriptional pattern to connect them to the mating process, but the mating pattern is not in conflict with that hypothesis. The balance of DedA and SNARE homologs may be mediating the fusion required for mating and are considered a primary target for future work. Additionally, genes implicated in mediating cell shape (the *cetZ* genes) may potentially be involved in formation of the cytoplasmic bridge and many of those genes have a pattern of expression consistent with mating warranting further examination.

This work also identified significantly increased expression under a mating condition for selfish genetic elements and host defense genes. The increased expression of insertion sequence transposases indicates that mating conditions may be activating transposition, a finding reminiscent of the stress induced transposition burst of an IS element in *Halobacterium halobium* ^109^. Simultaneously, expression of the host defense, CRISPR-Cas and RM Systems, are also activated under mating conditions, likely limiting recombination and defending against non-self, invading DNA, and suggesting post-fusion barriers against mating are present in *Hfx. volcanii*.

This study has identified a number of genes in addition to those discussed that follow an overall pattern of expression consistent with mating efficiency observations. Many of those genes are of unknown function and would benefit from further computational and experimental analysis to identify potential links to mating or to other important aspects of *Haloferax volcanii* molecular processes. Work on this intriguing method of DNA transfer remains ongoing.

## Acknowledgements

The Computational Biology Core, Institute for Systems Genomics, University of Connecticut provided computational resources.

This work was supported through grants from the Binational Science Foundation (BSF 2013061); the National Science Foundation (NSF/MCB 1716046) within the BSF-NSF joint research program and NASA Exobiology (NNX15AM09G, and 80NSSC18K1533).

## Author contributions

RTP and AMM designed experiments and JPG and ASL designed computational analysis. AMM carried out experiments. RTP, JPG, ASL, AMM, NRM, and UG interpreted the analytical output. AMM, ASL, RTP, and JPG wrote the manuscript. ASL prepared the figures and tables. RTP, JPG, AMM, ASL, NRM, and UG reviewed and approved the manuscript.

