## Supplemental figures and note for "Insights into gene expression changes under conditions that facilitate horizontal gene transfer (mating) of a model Archaeon"

### Supplementary Note
### Over and under expressed genes

Differentially expressed genes were ordered by fold change for each time point to identify the genes that stand out (Supplementary Fig. S7). While a number of genes were found to have higher expression than the shaking control at multiple time points, all but six of the genes with lower expression were only represented at one time point. The 2h time point had high differential expression of multiple ribosomal proteins, expression that was not observed in any other time point. HVO_RS19620, over expressed at 2h, may be a small CPxCG-related zinc finger protein, as per the European Nucleotide Archive annotation (ENA). Another putative zinc finger protein (HVO_RS01045) currently annotated as hypothetical was in the top ten highest fold change genes for all five time points. Recently, a similar set of zinc finger proteins was examined for their role in stress adaptation and biofilm formation (Nagel et al., 2019). One gene, HVO_RS07900, is annotated as ribose-1,5-bisphosphate isomerase, but has since been reannotated as thiazole synthase Thi4 (Hwang et al., 2014). Similarly, HVO_RS00195 is annotated as an iron transporter, but has been updated to be siderophore biosynthesis protein IucC and HVO_RS00200 has been identified as IucD (Niessen and Soppa, 2020). Many genes are annotated as ABC transporter subunits and highly expressed. HVO_RS16135 may be a putative phosphate transporter, HVO_RS04590 may be a putative zinc transporter, and HVO_RS00975 and HVO_RS12900 may be putative iron transporters.

Many of the genes that had the highest fold changes at each time point were listed as “hypothetical proteins” without a known or putative function. A homology analysis revealed potential functions for some of those genes. For example, HVO_RS04575 is globally homologous (probability: 99.2) to a heme-based aerotactic transducer HemAT found in *Haloferax* sp. BAB2207. HVO_RS20110, over expressed at 0 and 4-hours, may also be a small CPxCG-related zinc finger protein and has homologs in a large number of other archaea, as well as in bacteria, with high statistical significance (probability > 95, p-value < $1.1e^{-8}$). HVO_RS12600 is homologous to the caa(3)-type oxidase, subunit IV in *Desulfovibrio desulfuricans* ND132 (probability: 99.8) and is over expressed at 4 and 8 hours. HVO_RS02110, over expressed at 8h, is a homolog to the glycoprotein 13 gene (probability: 99.4). Also over expressed at 8h was HVO_RS06550, homologous to an RNA polymerase ECF-subfamily sigma factor in *Streptomyces filamentosus* (probability: 99.7) and similar proteins in dozens of organisms at >99% probability.

Some of the hypothetical proteins examined did not have a significant match to any gene in the database with greater than 95% probability. For example, the nearest homologous hits to HVO_RS19890, over expressed at 0h and 24h, were two uncharacterized proteins in a bacterium and an archaeon and the third nearest hit was to an uncharacterized protein in the Tree Cotton genome (probabilities: 84.0, 82.4, and 80.7, respectively). Another gene, HVO_RS19895, over expressed at all except the 2h time point, had a few more distant homologous matches to other *Haloferax* spp. and may be a cytochrome oxidase with homology to that gene in a Betaproteobacteria (probability: 91.4). With greater than 99% probability, HVO_RS03635 over expressed at 4h and 24h, and HVO_RS02000, over expressed at 24h, each had homology in six or more other Haloarchaea, but the nearest functional assignment to both was to the leucine-rich repeat-containing protein 51 in a choanoflagellate aligning over only a 35 amino acid region (probability: 84.7 and 85.0, respectively). Similarly, HVO_RS17490, also over expressed at 4h, has its closest homology to the outer capsid protein VP4 in Bovine rotavirus A, but not with high significance (probability: 86.9) and a significant homolog in only one other Haloarchaea.

There were also a number of under expressed genes listed as hypothetical proteins. The under expressed genes had low gene expression compared to the shaking control. Many of those genes did not have a statistically significant homolog with the following exceptions. HVO_RS13770 was under expressed at 0 hours and had high homology to a PINc domain-containing protein in Desulfuromonas soudanensis (probability: 96.2). One gene under expressed at 2h, HVO_RS11000, is a statistically significant homolog to the type I phosphodiesterase/nucleotide pyrophosphatase found in *Haloferax* sp. BAB2207 (probability: 100.0). Also under expressed at the 2-hour time point were HVO_RS08795, a homolog to a HxlR family transcriptional regulator (probability: 97.6) and HVO_RS08885, homologous to an uncharacterized archaeal protein with a potential isoform iron-sulfur cluster carrier protein (probability: 100.0). HVO_RS07195 is homologous to a zinc finger found in FPG and IleRS from bacteria (probability: 94.5) and under expressed at 4 hours. HVO_RS14495, under expressed at 8 hours, was homologous to the *Peptococcaceae bacterium* SHOCT domain-containing protein (probability: 99.7). Also under expressed at 8 hours was HVO_RS11890, homologous to an archaeal Arabinose efflux permease family protein (probability: 99.6). HVO_RS07070 was also under expressed at 8 hours and homologous to an FmdB family regulatory protein (probability: 99.5). While not a statistically high probability, HVO_RS02490 is homologous to a Glycosyltransferase family protein from *Haloferax* sp. BAB2207 (probability: 88.8). Under expressed at 24 hours, HVO_RS15670 is homologous to *Haloquadratum walsbyi* F pilus assembly type-IV secretion system for plasmid transfer (probability: 99.8).

The homologous gene search for hypothetical proteins also revealed potential links to N-glycosylation, which were examine because this pathway has experimentally confirmed links to mating and this transcriptome analysis has identified support for additional links. HVO_RS20230, HVO_RS02745, and HVO_RS16215 are homologous to putative membrane proteins required for N-linked glycosylation two in *Methanobacterium* and one a halophilic archaeon, respectively. HVO_RS15595 is homologous to *pglG*, an N-linked glycosylation glycosyltransferase in *Pseudoalteromonas* and was highly statistically significant at 2, 4, and 8 hours ($p$-value $< 0.01$) and significant at 24 hours ($p$-value $< 0.02$). HVO_RS02695 is homologous to *pglB*, pilin glycosylation protein, in *Neisseria meningitidis*. HVO_RS02800 is homologous to a general glycosylation pathway protein in *Campylobacter upsaliensis* and was differential expression was statistically significant at 8 hours ($p$-value $< 0.05$). HVO_RS08080 is homologous to a putative membrane oligosaccharyl transferase required for N-linked glycosylation in *Methanocella conradii*. The expression patterns for two of these genes, HVO_RS02800 and HVO_RS15595, followed the mating pattern and were the only two with statistical significance.

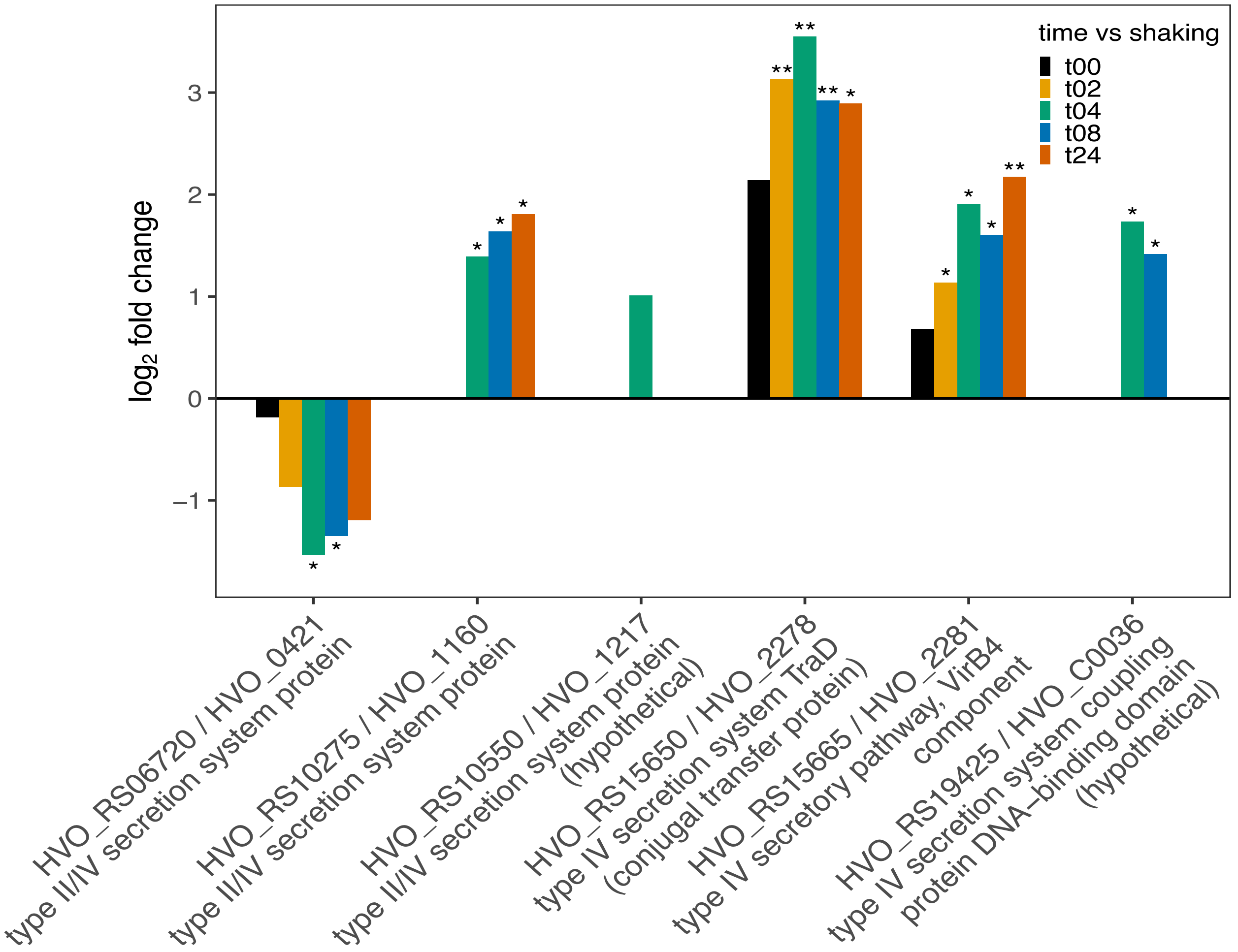


Supplemental Figure S1: Bar plot depiction of differential expression compared to planktonic state for type IV secretion system genes; gene id on the x-axis and log2 fold change on the y-axis, *p-value <0.05 and **p-value<0.01.

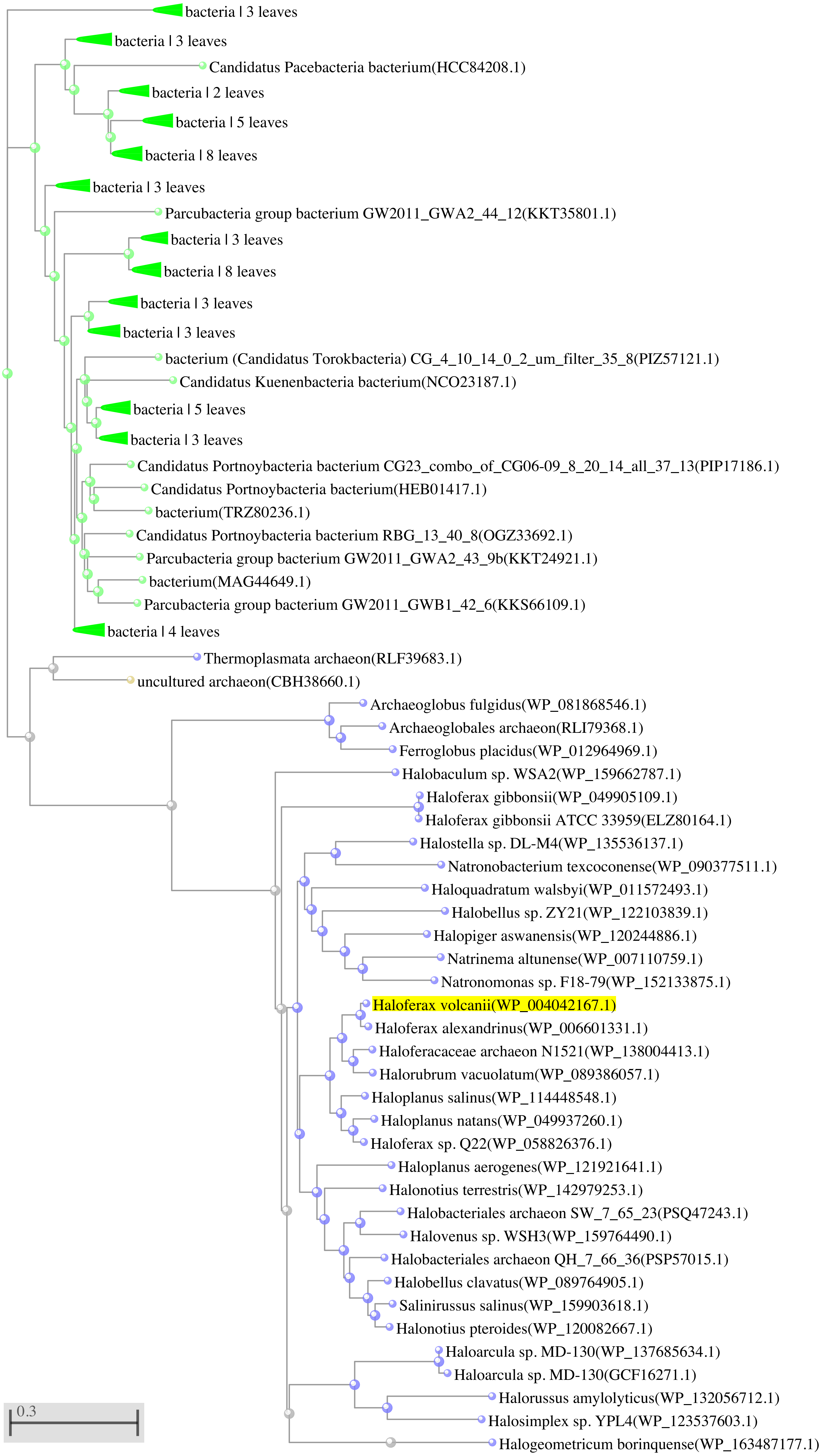


Supplemental Figure S2: Phylogeny of homologs to HVO_RS15650, conjugal transfer protein, across both Archaeal and Bacterial domains.

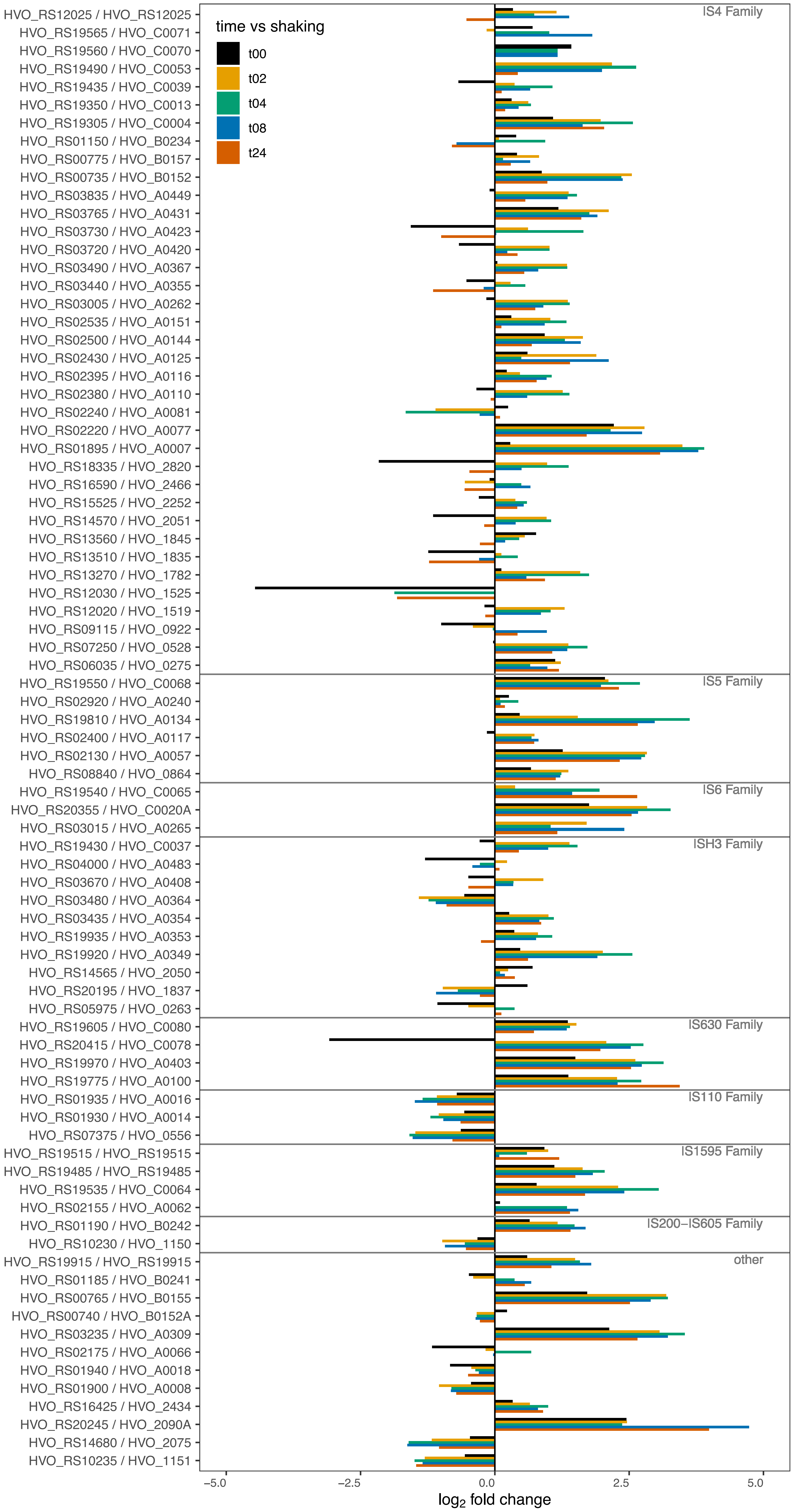


Supplemental Figure S3. Bar plot depiction of differential expression compared to planktonic state for insertion sequence gene families; gene id on the y-axis and log2 fold change on the x-axis.

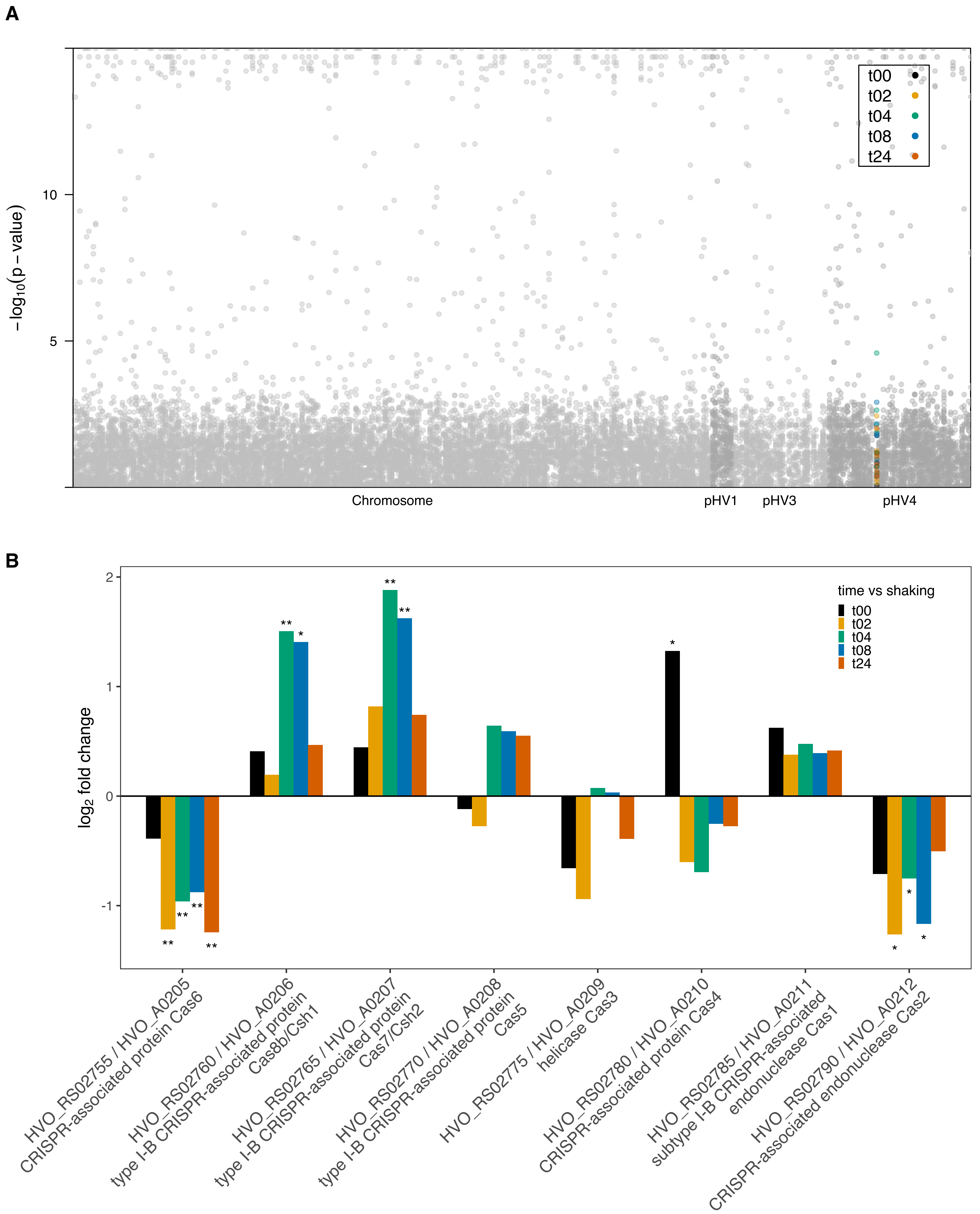


Supplemental Figure S4. (A) Manhattan plot with chromosome number on the x-axis and -log10 p-value of differential expression for planktonic cells vs plated timepoints on the y-axis, all genes are plotted in grey and genes for CRISPR-Cas are highlighted by color according to the key; (B) Bar plot depiction of differential expression compared to planktonic state for CRISPR-Cas genes; gene id on the x-axis and log2 fold change on the y-axis, *p-value <0.05 and **p-value<0.01.

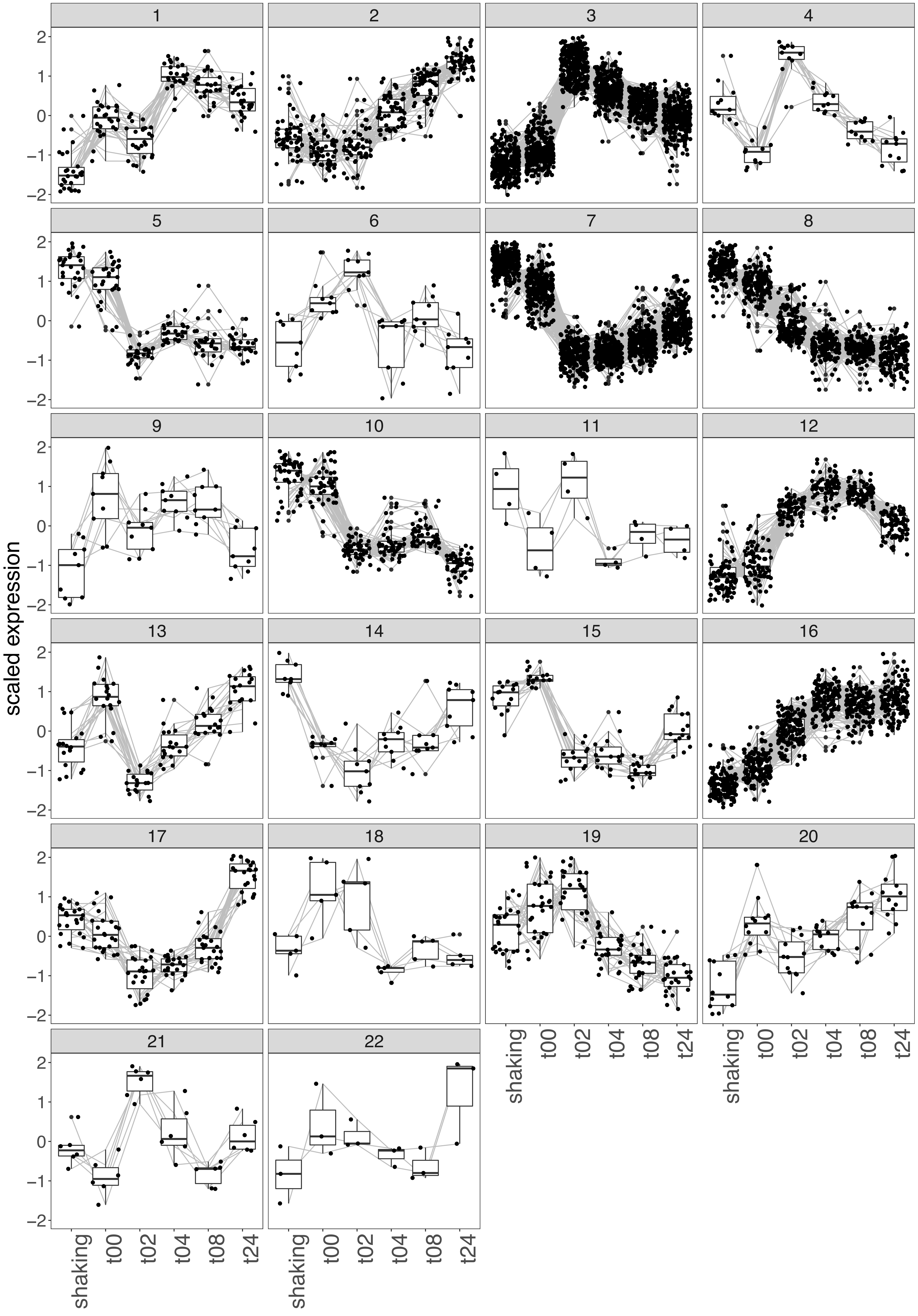


Supplemental Figure S5: Clustering of all expressed Haloferax volcanii genes for identification of similar expression patterns; time is on the x-axis and expression values scaled to the mean are on the y-axis.

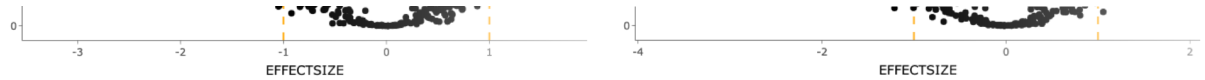


Supplemental Figure S6: Volcano plots of differential expression for the genes in cluster three comparing shaking to each time-point as labeled; x-axis is log2 fold change, y-axis is -log10 p-value, orange lines mark log2 fold change of -1 and 1, and green line marks a p-value of 0.01. .

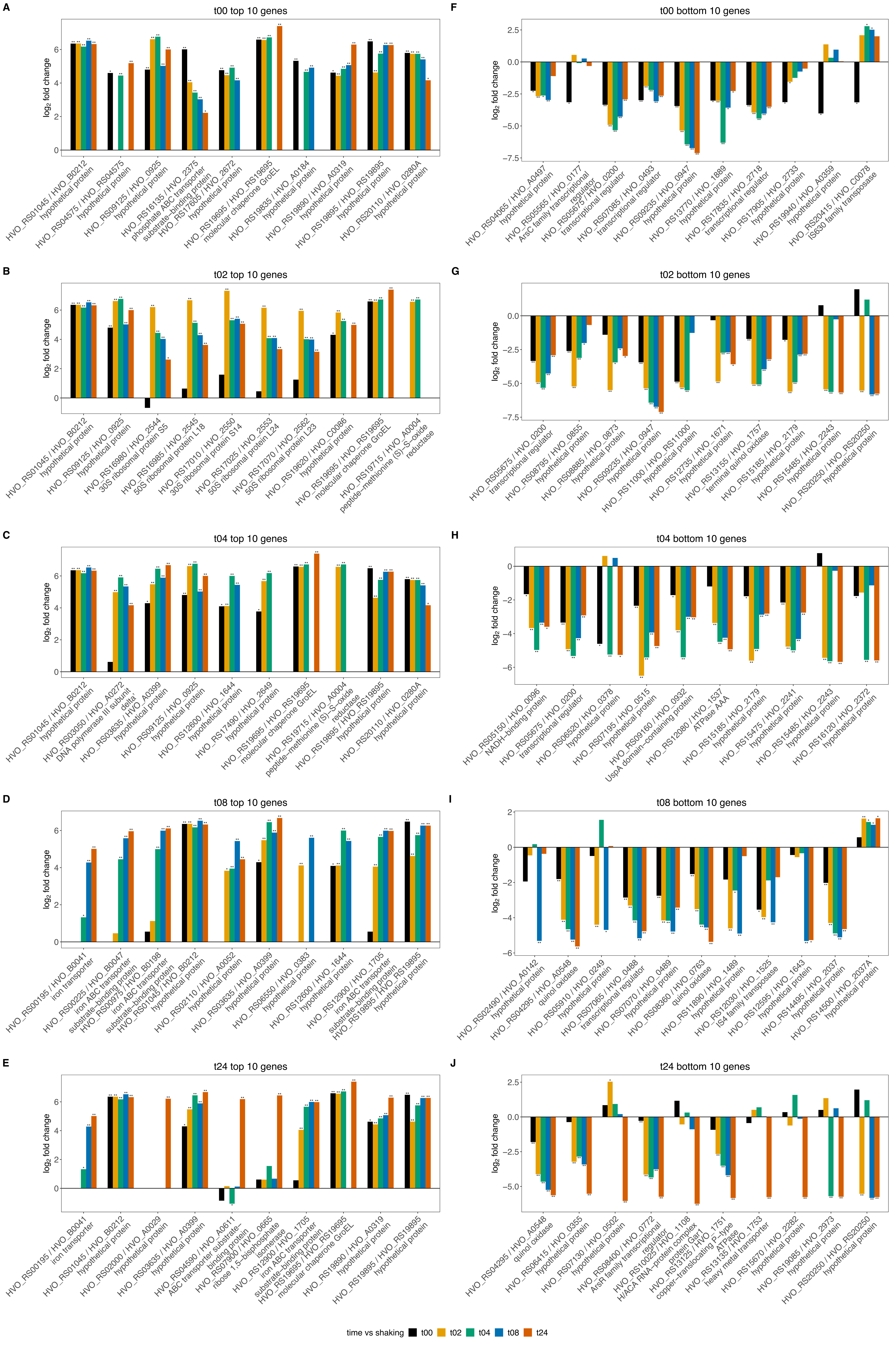


Supplemental Figure S7:Bar plot depiction of differential expression compared to planktonic state for top and bottom ten genes for each time-point; gene id on the x-axis, log2 fold change on the y-axis, and titles indicating the time-point and top or bottom; *p-value <0.05 and **p-value<0.01.

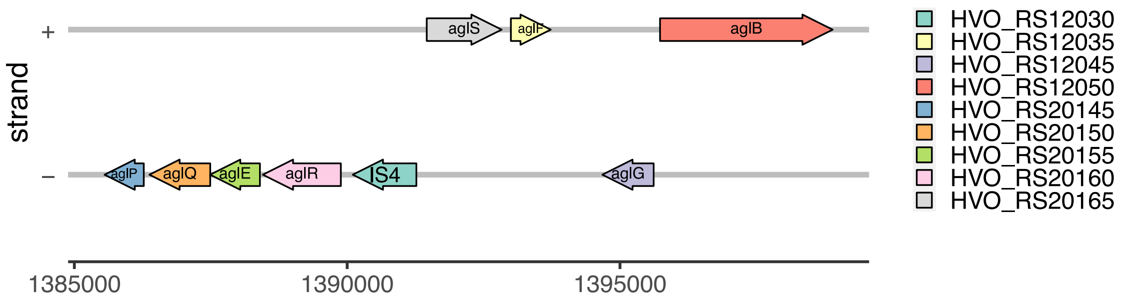


Supplemental Figure S8: Gene map of co-located glycosylation genes in cluster 3.
